# Adjuvants to improve efficacy of miticides in managed honey bee (*Apis mellifera*) colonies to control *Varroa destructor*

**DOI:** 10.1101/2025.02.14.638256

**Authors:** Brandon Shannon, Rui Zhang, Lucas Marsh, Reed M. Johnson

## Abstract

Beekeepers must manage *Varroa destructor* mites to maintain colony health. Large-scale beekeepers often use chemical treatments (miticides) to manage this pest. Miticide resistance drives a need for compounds with alternative modes of toxic action that can be used in a rotation as part of a *Varroa* management plan. This research aimed to determine the efficacy of oxalic acid, clove oil, and fenpyroximate when delivered in glycerin soaked in strips and combined with a range of bee-safe adjuvants. Adjuvants are a group of compounds used in plant pesticide applications to increase the spreading and penetration of a pesticide. Laboratory cage trials tested a miticidal active ingredient (oxalic acid, clove oil, or fenpyroximate) and an adjuvant (Ecostep BC-12^®^, Ecostep SE-11^®^, Ecostep AE-13^®^, Ecostep CE-13^®^, or Silwet L-7500^®^) in glycerin-soaked strips; field trials evaluated oxalic acid combined with Ecostep BC-12^®^ adjuvant in glycerin-soaked strips. Field trials with oxalic acid and adjuvant in glycerin caused a significant decrease in *Varroa* in year 1 and a significant reduction in *Varroa*, relative to the solvent control, in year 2. On the other hand, field trials with oxalic acid alone did not reduce *Varroa* loads in colonies relative to the solvent control. Additionally, mite drop data indicate increased speed for the miticidal effect when an adjuvant is included with oxalic acid. This research informs formulation chemistries for oxalic acid and other miticides to help beekeepers maintain healthy hives.

## Introduction

Honey bees (*Apis mellifera*) are responsible for pollination of at least 100 crops (1–3), with an estimated contribution of 12 billion dollars to the US economy (4,5) and 153 billion USD worldwide (6). However, high colony losses have been reported by commercial beekeepers, at an average annual colony loss rate of 55% and winter loss rate of 37% in the United States (7) and an average winter loss rate of 34% in Canada in 2023-2024 (8). Parasitism by the ectoparasitic mite *Varroa destructor* is a major driver of colony loss. In surveys beekeepers self-reported that *Varroa* and its associated diseases were the cause of about 60% of winter colony losses in U.S. commercial beekeeping operations (9,10). Pathogens carried by *Varroa* including deformed wing viruses, black queen cell virus, acute paralysis viruses, and nosema, among others (11–18).

Because many of the non-chemical management strategies for *Varroa* are not sufficiently effective or too labor-intensive (19,20), large-scale beekeepers must use chemical treatments, known as miticides, to manage this pest. However, over-reliance and misuse of synthetic miticides have led *Varroa* to develop worldwide resistance to the formamidine amitraz (21–26), the organophosphate coumaphos (27–30), and the pyrethroids fluvalinate and flumethrin (31–34). Because of widespread miticide resistance, there is a rising need for products with alternative modes of toxic action that can be used in a rotation as part of a *Varroa* management plan. Potential alternatives explored by this study include extended-release formulations of the natural pesticides oxalic acid and clove oil, and the synthetic pyrazole fenpyroximate.

Oxalic acid is a miticide that is permitted for use in the United States, several European countries, and New Zealand (20), and has been used to control *Varroa* for several decades (35). The mechanism of toxic action of oxalic acid to *Varroa* is not well understood (36), though effects of physical damage to the chitin plate of *Varroa* have been observed (37). However, oxalic acid is not effective against *Varroa* protected by brood cell cappings, so it has limited efficacy when applied as a vaporization or dribble flash treatment (38,39). At the time of this study, no oxalic acid extended-release treatments were available for use in the United States, but many beekeepers treated with homemade oxalic acid extended release formulations using sponges or shop towels soaked in a 1:1 w/w mixture of glycerin and oxalic acid (40). While clove oil (principal constituent eugenol) does not have an defined IRAC mode of action, it has demonstrated contact toxicity to *Varroa* (41–44). Clove oil can act as an insect repellent and prohibits the growth of bacteria and fungi (45), but likely affects *Varroa* through effects on the enzymes glutathione-S-transferase, superoxide dismutase, and Ca^2+^-Mg^2+^-ATPase (46). Fenpyroximate is a synthetic pyrazole acaricide with an IRAC class 21A Mitochrondrial Complex I Electron Transport Inhibitor mode of toxic action (47) that was previously registered for in-hive use as Hivastan® and demonstrates promising mite control in semi-field evaluations (48). Oxalic acid is registered for use as a dribble or vaporization treatment as Api-Bioxal®, as a vaporization treatment as EZ Ox®, or as an extended-release strip formulation as Varroxsan® (49), though only Api-Bioxal® was registered for use at the start of this experiment. Neither clove oil nor fenpyroximate are currently formulated in registered *Varroa* control products.

Adjuvants are added to pesticides, either as formulation components or tank-mix components, to improve the handling or application characteristics and enhance pesticide activity (50). While adjuvants are typically used with spray applications in agriculture, they may also be included in some pesticides used to control common bee pests in managed honey bee colonies (51), though the identity of adjuvant ingredients in these products are proprietary. The “principal functioning agents” that provide the desired function of an adjuvant are drawn from the list of inert ingredients maintained by US EPA and consist of the same or similar compounds used as formulation components in traditional pesticide products (52). While some adjuvants have shown high toxicity to honey bees, others are relatively non-toxic (51,53). Using these adjuvants that demonstrate low toxicity to bees as formulation components in *Varroa* control applications can potentially improve miticide activity to and greaten the number of effective active ingredients for control of *Varroa* mites (54). (52)

Oxalic acid, clove oil, and fenpyroximate, have shown promise as miticides for *Varroa* control, but demonstrate limited efficacy at concentrations that are safe for honey bees and none are known to be effective at controlling *Varroa* on pupae that are protected by brood cell cappings. This research aimed to determine the efficacy of these active ingredients when delivered in glycerin soaked in strips and combined with a range of bee-safe adjuvants. Combinations of active ingredients and adjuvant were first tested in laboratory cage trials to identify the most promising miticide-adjuvant combination, which was then assessed in a field trial to determine if the addition of an adjuvant could improve varroa control in whole honey bee colonies.

## Materials and Methods

### Miticides and Adjuvants

The miticides used in the laboratory cage trial included clove oil (100%; Sigma-Aldrich; MO, USA); fenpyroximate (>98.0%; Alfa Chemistry, Inc.; NY, USA); and oxalic acid dihydrate (99.5 – 102.5%; Thermo Fisher Sceintific; MA, USA). The miticide used in the field trial was oxalic acid dihydrate.

The adjuvants used in the laboratory cage trials are listed in Table 1 and include Ecostep AE-13® (Stepan, Northfield, IL, USA), Ecostep BC-12® (Stepan), Ecostep CE-13® (Stepan), Ecostep SE-11® (Stepan), and Silwet L-7500 Copolymer® (Momentive Performance Materials, Niskayuna, NY, USA). The adjuvant used in the field trial was Ecostep BC-12®.

**Table 1.** Adjuvant treatments used in this experiment. The principal functioning agents for all adjuvants are included in the list of inert ingredients safe for food and nonfood use (52). An asterisk (*) indicates that the concentration of principal functioning agents is not given and the SDS states that the components are not hazardous or are below required disclosure limits.

| Treatment | Manufacturer | Principal Functioning Agents (% Composition) |
| --- | --- | --- |
| Ecotstep AE-13® | Stepan | Polyethylene Glycol / Polypropylene Glycol - (n/n) Copolymer (> 99%) |
| Ecotstep BC-12® | Stepan | Polyethylene / Polypropylene Glycol Monobutyl Ether (90 – 100%) |
| Ecotstep CE-13® | Stepan | Castor Oil, Ethoxylated (90 – 100%) |
| Ecotstep SE-11® | Stepan | Sorbitan, Mono-(9z)-octadecenoate, Poly(oxy-1,2-ethanediyl) Derivs (90 – 100%) |
| Silwet L-7500 Copolymer® | Momentive Performance Materials | Polyalkyleneoxide modified polydimethylsiloxane* |

### Adjuvant acute toxicity tests

All adjuvants were tested for contact bee toxicity using a Potter Spray Tower (55). The insecticide Mustang Maxx ^®^ (9.15% zeta-cypermethrin active ingredient), a pyrethroid insecticide with high acute toxicity to honey bees, was used as a positive control in the Potter Spray Tower toxicity tests (53,56–58). The 48-hour acute contact LC_50_ of all adjuvants were determined using 3-day-old adult worker bees following methods described in (53). The concentrations tested were as follows: for the four Ecostep® products, 0, 1, 3, 5, 10, and 20% by volume; for Silwet L-7500 Copolymer®, 0, 0.3, 1, 3, 5, 10, and 20% by volume; for Mustang Maxx® Positive Control, 0.00186, 0.00558, 0.0186, 0.0558, and 0.186% by volume of formulation, which is 1.70e-4, 5.10e-4, 1.70e-3, 5.10e-3, 1.70e-2% zeta-cypermethrin active ingredient by volume (53,58).

### Cage Trials

#### Treatment Preparation

Treatment solutions were prepared by adding glycerin (Fisher Scientific; MA, USA) to a 15 mL conical tube (Thermo Fisher Scientific; MA, USA) and sonication with heating (Kendal Digital Ultrasonic Heated Cleaner model HB-S-49DHT) for 30 min to make 3 g of final treatment mixture. Negative control treatment solutions containing only glycerin were also heated. Active ingredient concentrations were based on preliminary cage trials that measured bee mortality. Clove oil treatments were prepared by adding clove oil to form a 3% v/w mixture with heating at 34° C. Fenpyroximate treatments were prepared by first dissolving solid fenpyroximate in acetone to a 10% w/v solution, then adding the fenpyroximate-acetone solution to glycerin to produce a 0.2% w/w solution that was heated to 34° C. Oxalic acid treatments were prepared by adding oxalic acid dihydrate to glycerin, up to, but not above, 67° C (59–62) and sonication for a minimum of 30 minutes, until crystals were no longer visible, to produce a 20% w/w solution. The concentrations of the active ingredients in cage trials were chosen based on preliminary studies that determined the maximum concentrations that could be applied with minimal mortality observed in caged bees.

Active ingredient – adjuvant combination treatments were prepared by adding a single adjuvant product (Ecostep AE-13®, Ecostep BC-12®, Ecostep CE-13®, Ecostep SE-11®, or Silwet L-7500 Copolymer®) to the glycerin-active ingredient mixture and mixed fully to produce a 0.5% w/w solution for the Ecostep products or a 0.2% w/w solution for Silwet L-7500 Copolymer. The final active ingredient concentration was the same as in the active ingredient controls. The concentration of adjuvant was chosen based on the labeled concentrations for similar adjuvants in agricultural tank mix applications, which typically range from 0.0625 to 0.625 percent by volume (53).

Cage treatment strips were made from a Swedish sponge (Superscandi Swedish Dishcloths; London, England) cut into 32 sections (1.25 x 8.5 cm). A single treatment strip was placed in each 3 g solution and heated with sonication at the respective temperature for each treatment for 30 minutes. The concentration of 1/32 saturated treatment strip was used in cages to scale down a full-size strip that would be used with 10,000 bees in a deep Langstroth box (63,64) to approximately 300 bees in each cage.

#### Cage Design

Cage design was modified from (Rinkevich, 2020; S1 Text, Fig A). The 1.25 x 8.5 cm treatment strip was inserted in a 15 mm x 3 mm slit cut in the top of the cage. Four equally spaced 3-mm holes were added for airflow on the sides of the cage, 1 cm from the top of the cage. Two 3-5 g sucrose cubes were fixed to the top of the cage using hot melt adhesive to allow bees to feed.

#### Honey Bees

To populate the cages, honey bees were shaken from brood frames from six colonies, in July through October in 2022 and 2023. Colonies were managed at The Ohio State University – Wooster campus apiaries and had not been treated for mites for at least 6 months prior to collection. All colonies had been requeened with New World Carniolan (Olivarez Honey Bees, Orland, CA) in the Spring, but queens were caged for at least 21 days prior to worker bee collection to maximize the number of phoretic mites. Bees were shaken into an empty 4-frame nucleus box and stored in darkness for no longer than one hour before transfer to cages. Bees were misted with DI water during transfer to discourage flight. Approximately 300 bees were scooped from the nucleus box and placed into each cage containing a treatment.

#### Experimental Design

Each cage treatment series consisted of a negative control, an active ingredient control, and four active ingredient-adjuvant combinations. Treatments were assigned in random order. Cages were placed inside an incubator (Humidaire Model No. 2048; The Humidaire Incubator Company, New Madison, OH, USA) and stored at hive conditions (34° C, 60% humidity, darkness). Dead bees at the bottom of the cage and dead *Varroa* on the collection tray were counted after 24 hours. After bees and *Varroa* were counted, the plastic tray was discarded and cups were inverted and frozen at -20° C.

Bees were weighed (Ohaus, Parsippany, NJ; Model CL5000) and the number of grams was multiplied by 11.34 to estimate the number of bees, as determined in preliminary trials. Bees were agitated for 30 minutes in 70% ethanol to dislodge *Varroa* remaining on the bees for counting. Treatment efficacy was determined by dividing the number of *Varroa* that had fallen during treatment by the total number of *Varroa* (mites fallen plus mites in alcohol wash). A Kruskal-Wallace test was used to determine significant differences in bee mortality and *Varroa* control efficacy. This was followed with a pairwise Wilcoxan Rank-Sum Test using Benjamini-Hochberg post-hoc correction.

### Field Trials

#### Honey bee colonies

Three apiaries, separated by a minimum of 5 km, located at The Ohio State University – Wooster campus, were used to conduct the field trial. Each colony consisted of a minimum of two deep 8-frame Langstroth boxes at the start of the experiment. In year 1, each colony was fitted with a screened bottom board for collecting mite drop data. In year 1, 9 hives from apiary 1, and 6 hives each from apiaries 2 and 3 (21 total) were randomly assigned treatments. In year 2, 9 hives from each of the 3 apiaries (27 total) were randomly assigned treatments. In both years, treatments were stratified based on pre-treatment *Varroa* levels and were assigned so that each apiary had an equal number of replicates of each treatment.

#### Experimental Design

Before and after treatment, colonies were assessed by performing *Varroa* alcohol washes and seam counts (65–67). Seam counts are used to estimate the number of adult bees in a colony and involves two observers visually estimating the clusters or “seams” of bees found between the frames in each box of the hive. Seams in medium boxes were multiplied by the height of a medium Langstroth box divided by the height of a deep Langstroth box (6.625/9.625). Any colonies with less than 9 seams prior to treatment or less than 2 seams after the treatment were excluded from analysis. Alcohol washes were performed by collecting approximately 300 bees from 3 worker brood frames into 70% ethanol, followed by shaking 30 minutes, and counting both bees and *Varroa* that were strained from the wash.

For year 1, *Varroa* that fell below the colony during the treatment, or mite drop, was monitored at 48-hour intervals starting 2 days prior to treatment, at the time of treatment application (day 0), and then 2, 4, 7, 14, and 21 days following treatment. Mite drop was monitored by placing letter-sized manilla file folders (Staples, Framingham, MA, USA) coated with Vaseline® petroleum jelly under screened bottom boards. After folders were removed, they were frozen at -20° C for a minimum of 24 hours and stored under these conditions until counted.

#### Treatments

In year 1, the adjuvant combination treatment consisted of 1% w/w Ecostep BC-12® adjuvant and 40% w/w oxalic acid dihydrate dissolved in glycerin, the oxalic acid control consisted of 40% w/w oxalic acid dihydrate dissolved in glycerin, and the glycerin control consisted of glycerin only. In year 2, a 0.5% w/w adjuvant concentration was used instead of the 1% concentration in the oxalic acid plus adjuvant treatment. For each uncut Swedish sponge, 70 g of solution, the amount needed for complete saturation of one sponge, was prepared and heated at less than 67° C for at least 1 hour with sonication (Kendal Digital Ultrasonic Heated Cleaner model HB-S-49DHT) and stirring, if necessary.

In year 1, one treatment strip, consisting of a full Swedish sponge (20.3 x 17.8 cm) saturated in treatment solution, was applied for every 9 seams of bees, rounded up, so that each colony had either 2 or 3 treatment strips. In year 2, strips were cut in half to increase contact area with bees, and strips were applied for every 5 seams of bees, rounded up, so that each colony had at least 3 treatment strips. Treatment strips were placed between boxes of brood and colonies were left undisturbed during the treatment period. In year 1, treatments were in place for 23 days starting on Sept 21st for apiary 1, Oct 9th for apiary 2, and Oct 14th for apiary 3. In year 2, treatments were applied for 22 days starting on July 30 for all apiaries.

#### Data Analysis

Field trial results were analyzed independently for both years, using the same statistical analysis methods. Efficacy within treatments was compared by performing a paired t-test on the difference between the pre- and post-treatment *Varroa* levels from alcohol washes using the stats package in R (68). The change in *Varroa* levels between treatments were compared by performing an ANOVA on the difference between pre- and post-treatment mite wash levels using the stats package in R (68). Mite drop for year 1 was analyzed using a Mixed Model for Repeated Measures (MMRM) on the cumulative mite drop data over the treatment period with the nlme and multcomp packages in R (69,70). Post-hoc comparisons were conducted on the cumulative mite drop to identify significant differences between treatments at specific time points (S2 File).

## Results

### Screening adjuvant toxicity to bees through a spray application

The 48-hour LC_50_ of all adjuvants tested was predicted to be greater than the maximum concentration sprayed with the Potter Tower (Table 2; S3 Dataset), and the mean mortality at the highest tested concentration for each adjuvant were below 50%. The LC_50_ for Mustang Maxx®, the positive control, was estimated to be 0.043%, (95% CI = 0.038, 0.047) when expressed as a formulation concentration, or 0.0039% (95% CI = 0.0035, 0.0044) when expressed as a concentration of zeta-cypermethrin active ingredient.

**Table 2.**
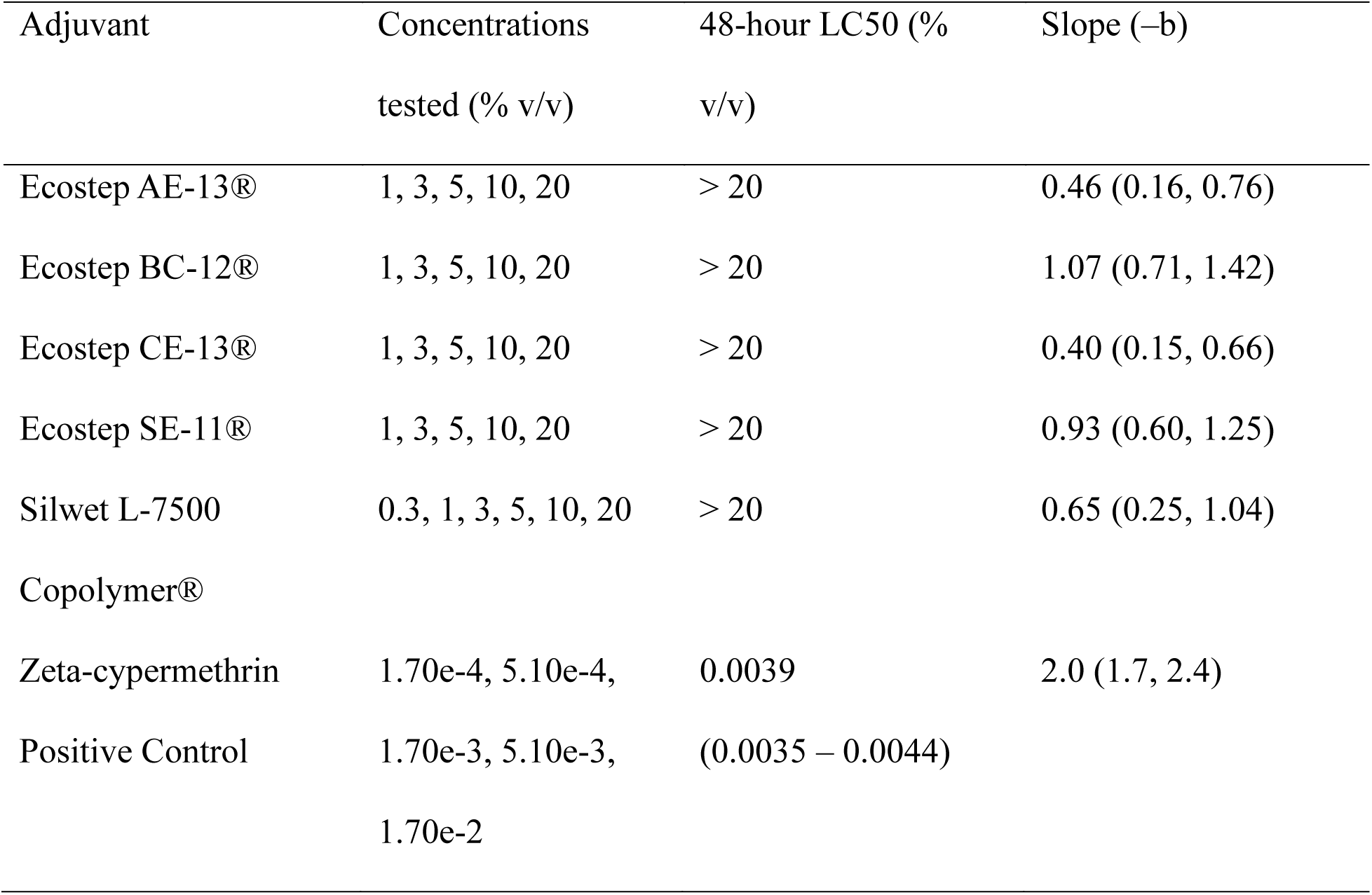
48-hour LC_50_ estimates of each adjuvant applied to adult bees using a Potter Spray Tower. All adjuvant LC_50_ estimates were above the maximum rate tested. Values in parenthesis indicate 95% confidence intervals.

### Cage Trials

#### Clove oil

There was no significant difference in 24-hour bee mortality among treatments (Kruskal-Wallace; P > 0.05; Fig 1A; S1 Text, Table A; S4 Dataset). A significant difference was observed in 24-hour *Varroa* efficacy (P = 0.0035; df = 5). However, a Pairwise Wilcoxon Rank-Sum Test using Benjamini-Hochberg post-hoc correction found no significant differences between treatments (P > 0.05).

**Fig 1.**
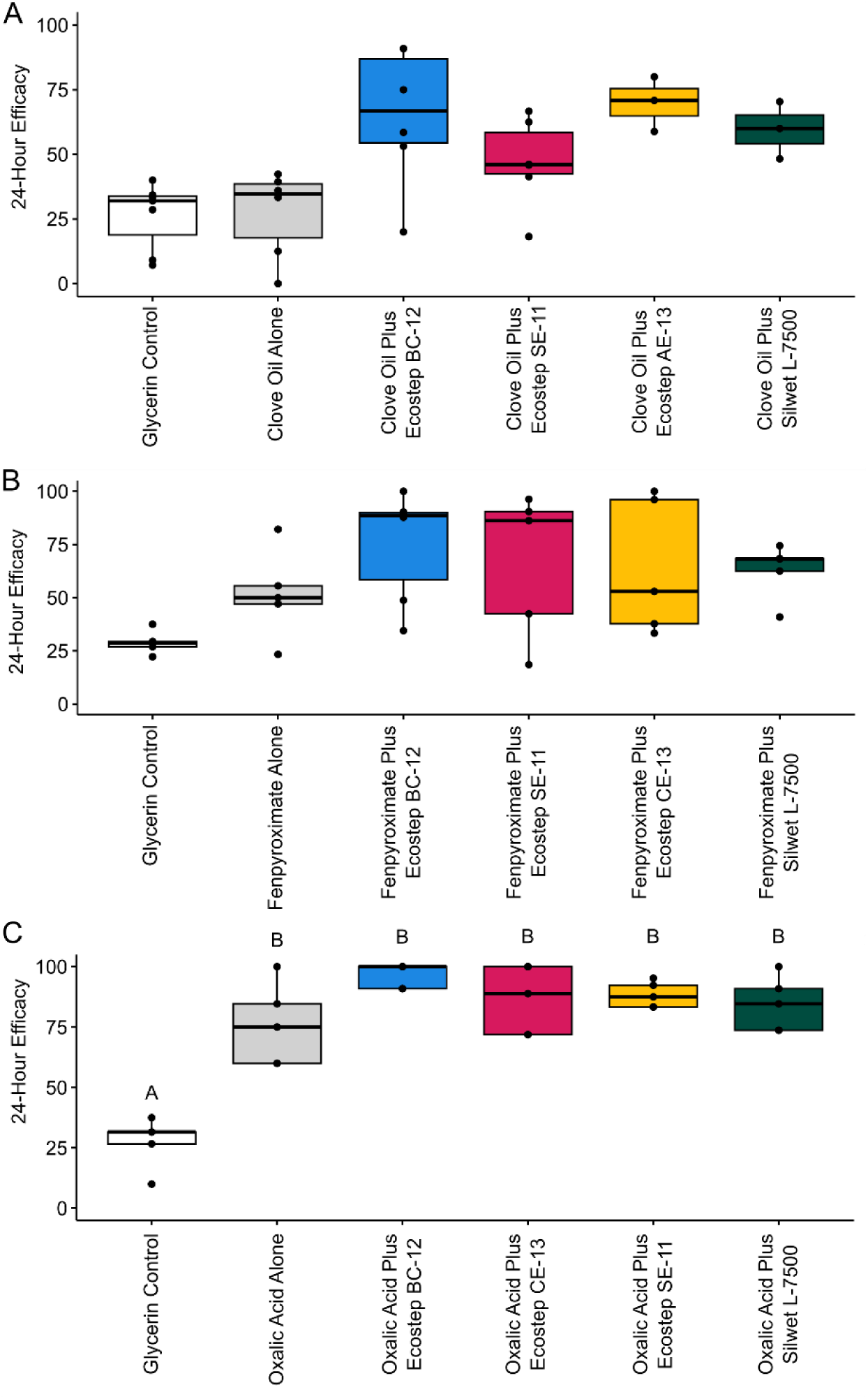
Efficacy of Clove Oil (A), Fenpyroximate (B), and Oxalic Acid (C) treatments in cage trials, where efficacy is determined to be the number of *Varroa* mites that fell in 24 hours divided by the total number of *Varroa* mites in each cage. There were no significant differences for clove oil or fenpyroximate treatments (P > 0.05). Different letters for oxalic acid treatments indicate significant differences (P < 0.05).

#### Fenpyroximate

There was a significant difference in 24-hour bee mortality (P = 0.034), but pairwise post-hoc tests determined no significant differences between treatments (P > 0.05; Fig 1B; S1 Text, Table A; S3 Dataset). There was no significant difference in 24-hour *Varroa* efficacy (P > 0.05).

#### Oxalic acid

There was no significant difference in 24-hour bee mortality between treatments (P = 0.34; Fig 1C; S1 Text, Table A; S3 Dataset). There was a significant difference in 24-hour *Varroa* efficacy (P = 0.0062). Pairwise post-hoc comparisons determined significant differences between negative control with active ingredient control and all oxalic acid-adjuvant combinations (P < 0.05), but no significant difference between the oxalic acid control and any oxalic acid-adjuvant combination (P > 0.05), though there was a trend for increased efficacy in all four adjuvant combinations.

### Field Trials

Colonies that had 2 or fewer seams of bees by the end of the experiment were excluded from analysis. In year 1, the analysis included 5 colonies from the glycerin control group, 7 colonies from the oxalic acid control group, and 6 colonies from the oxalic acid plus adjuvant treatment group. In year 2, no colonies were excluded, so all treatment groups included 9 colonies.

#### Pre- and Post-Treatment Ethanol Washes

In year 1, a significant decrease of 3.1 (95% CI = -4.1, -2.1) *Varroa* mites per 100 bees was observed between the post- and pre-treatment ethanol washes for the oxalic acid plus adjuvant treatment (one-sided t-test, P = 0.0122; n = 6, t = –3.189). The glycerin control had an increase of 8.4 (95% CI = 1.7, 15.1) and the oxalic acid alone had an average decrease of 4.3 (95% CI = -8.7, 0.1) mites per 100 bees, but neither demonstrated a significant difference between post- and pre-treatment ethanol washes (Fig 2A; S1 Text, Table B, Fig B; S5 Dataset). An ANOVA of the difference of the final and initial *Varroa* mites per 100 bees in ethanol washes determined no significant differences between treatments (P > 0.05).

**Fig 2.**
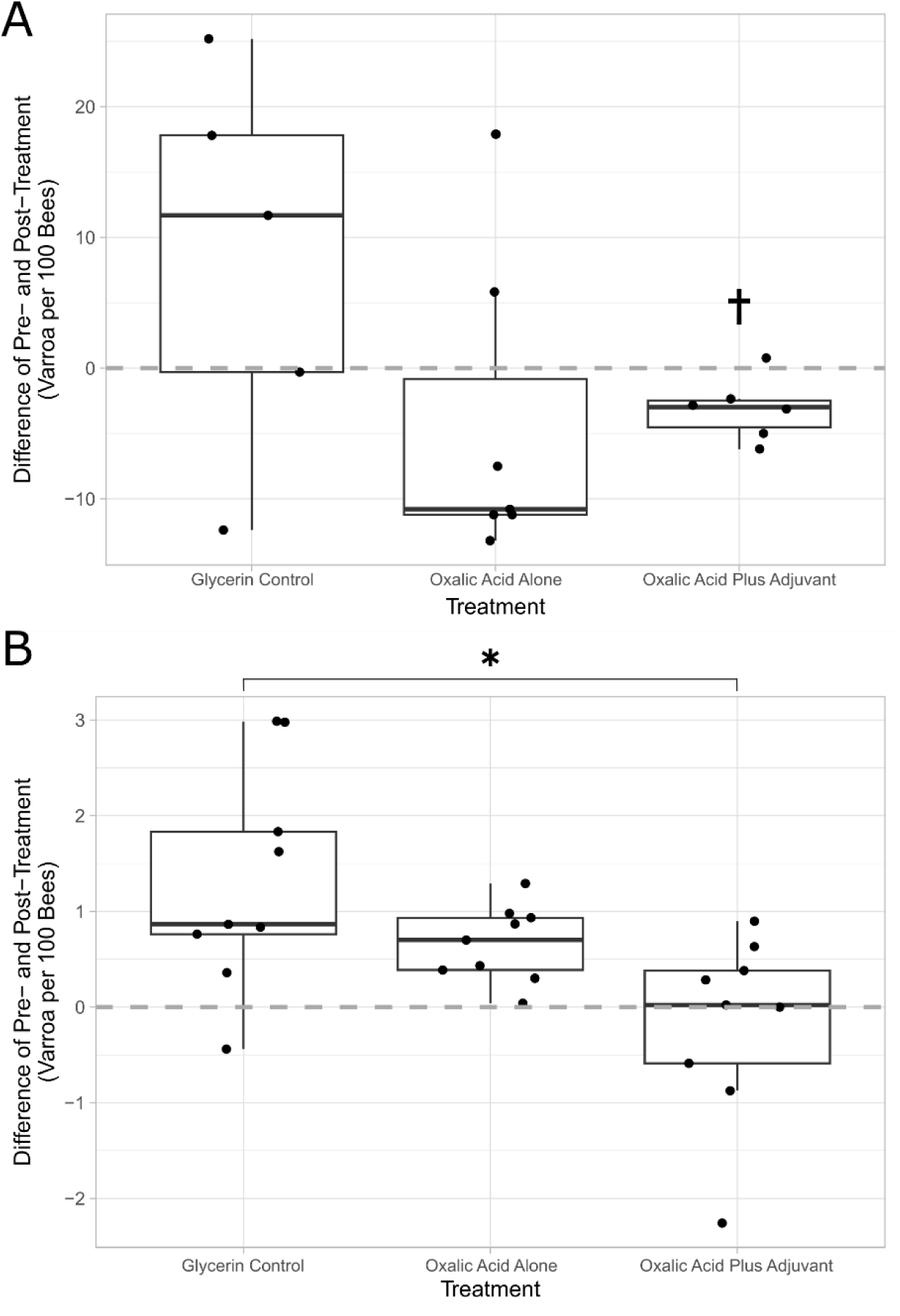
The change in *Varroa* levels from alcohol washes for year 1 (A) and year 2 (B) field trials. Points indicate the post-treatment minus the pre-treatment *Varroa* mites per 100 bees for individual colonies. A significant difference between treatments, indicated by an asterisk (*), was found between the glycerin control and oxalic acid plus adjuvant treatments in year 2 (P = 0.005). A significant difference between the pre- and post-*Varroa* mites per 100 bees, indicated by a dagger (†), was determined only for the oxalic acid plus adjuvant treatment in year 1 (one-sided t-test, P = 0.0122; n = 6, t = –3.189).

In year 2, the difference between the pre- and post-treatment *Varroa* in ethanol washes determined an average increase of 1.3 (95% CI = 0.9, 1.7) *Varroa* per 100 bees for the glycerin control, an average increase of 0.7 (95% CI = 0.5, 0.8) for the oxalic acid control, and an average decrease of 0.2 (95% CI = -0.5, 0.2) for the oxalic acid plus adjuvant treatment, but none were statistically significant (one-sided paired t-test, P > 0.05; Fig 2B; S1 Text, Table B, Fig B; S6 Dataset). An ANOVA between change in the initial and final *Varroa* from alcohol washes of treatments determined a significant difference between treatments (P = 0.007; df = 2, F = 6.143). A Tukey’s Post-Hoc test identified a significant difference between the glycerin control and the oxalic acid plus adjuvant treatment (P = 0.005), but no significant difference between the glycerin control and oxalic acid control or between the oxalic acid control and the oxalic acid plus adjuvant treatment (P > 0.05).

#### Varroa mite drop

There was a significant increase in *Varroa* mite drop over the first 4 days of treatment with oxalic acid plus adjuvant compared to oxalic acid alone (linear mixed-effects model; z = - 2.473, P = 0.0134), but only a marginal difference over the first six days of the treatment period (z = -1.74, P = 0.0819), and no significant difference for the full 23-day treatment period (z = - 1.182, P > 0.1; Table 3; Fig 3; S5 Dataset). Significantly more *Varroa* mites dropped in the oxalic acid plus adjuvant treatment relative to the glycerin control over the full 23-day treatment period (z = -4.35, P < 0.001). Similarly, comparing the oxalic acid alone to the glycerin control determined a significant difference for the full 23-day treatment period (z = 3.38, P < 0.001).

**Fig 3.**
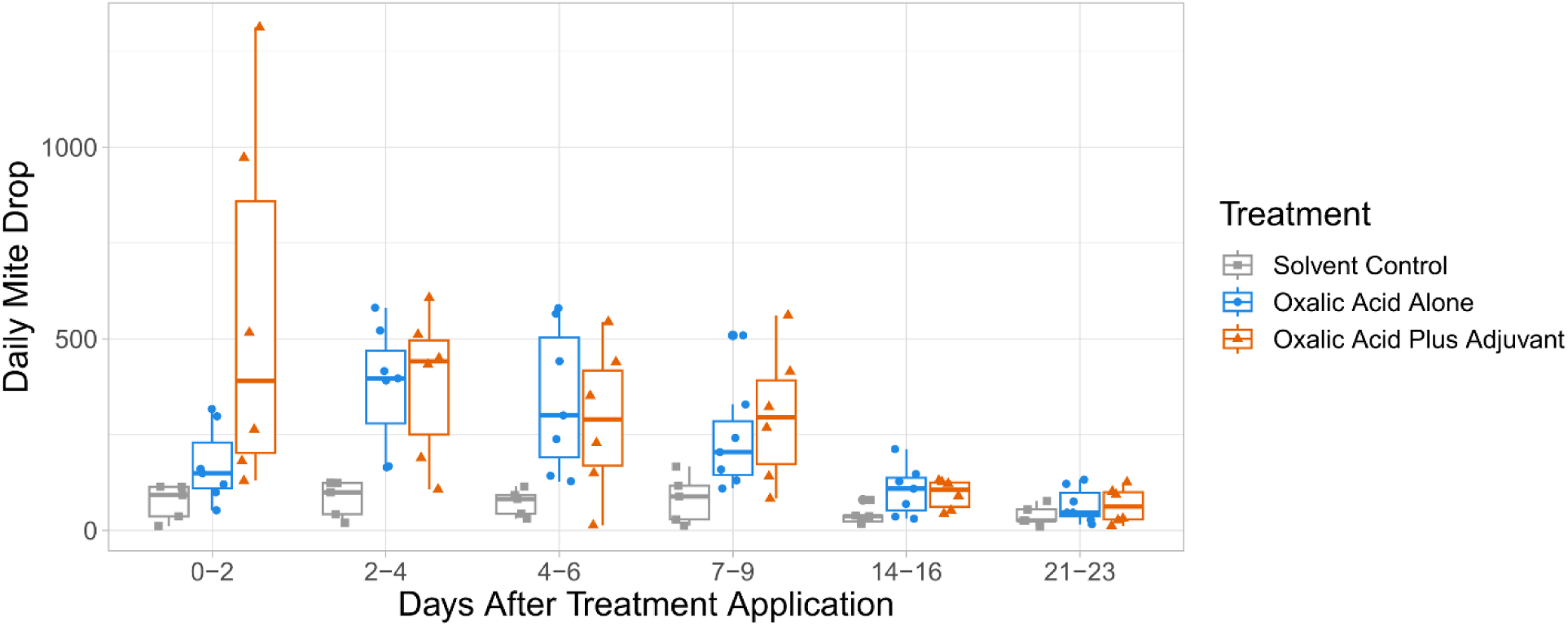
The daily mite drop for colonies in Year 1. Each point indicates the average daily mite drop over the 48-hour measurement period for an individual colony. Treatments were applied at day 0, immediately prior to sample collection.

**Table 3.** Average daily *Varroa* mite drop over the treatment period. Day 0 is the initial time of treatment. Baseline indicates daily mite drop for the 48-hour period prior to treatment application. Values indicate average daily mite drop during the measurement period for each treatment, with standard deviation listed in parenthesis, where baseline indicates the 48-hour period before treatment.

| Days after<br>treatment | Baseline | 0-2 | 2-4 | 4-6 | 7-9 | 14-16 | 21-23 |
| --- | --- | --- | --- | --- | --- | --- | --- |
| Glycerin<br>control | 103<br>(82.0) | 73.5<br>(47.1) | 81.9<br>(48.2) | 72.5<br>(34.8) | 82.4<br>(63.5) | 38.9<br>(24.5) | 38.4<br>(26.8) |
| Oxalic Acid<br>Control | 109<br>(72.1) | 171<br>(99.8) | 377<br>(160) | 342<br>(189) | 240<br>(140) | 104<br>(65.1) | 66.3<br>(45.4) |
| Oxalic Acid<br>Plus Adjuvant | 130<br>(76.8) | 563<br>(481) | 383<br>(194) | 288<br>(195) | 298<br>(176) | 94.0<br>(38.7) | 65.1<br>(47.4) |

## Discussion

Using a cage assay rather than a traditional topical application or glass vial application to *Varroa* better simulates the exposure that mites would face in a colony environment, but may introduce additional sources of variability compared to other topical application methods (71). Bees for cage trials were of unknown age, as they were collected by shaking frames. Therefore, there is variability between trails due to unequal age of bees, as age is a significant determinant of bee resilience to stressors (72–74), though age variability may provide greater field relevance. Additionally, different cohorts of bees have variability in *Varroa* loads, which have an impact on bee health (20,75,76) and low *Varroa* numbers can lead to an overrepresentation of efficacy against *Varroa* (71). Variable *Varroa* loads, and the effects on bee health that very high *Varroa* levels can have, were mitigated by excluding results from cages that were not within the range of 2 – 12 mites per 100 bees (71). The strength of the colony that bees were shaken from, especially as colony strength changes throughout the Fall, is an important factor in bee and *Varroa* resilience to stressors (77). This variability was managed by performing each set of treatments with the same cohort of shaken bees.

While the alcohol wash is often used as the standard procedure for *Varroa* detection in colonies, it is not without its faults (78,79). Alcohol washes using 70% ethanol in other studies have demonstrated 65 to 95% recovery of *Varroa* from the bodies of adult bees (80–82). However, this study used 30 minutes of rotational shaking, followed by 60 seconds of wrist-shaking, which exceeds the agitation times of each of the cited studies, so it is likely that the recovery of *Varroa* on adult bees in this study is near the highest cited value of 95%.

Cumulative mite drop rather than mite drop over individual 48-hour measurements was taken to avoid introducing a negative bias for treatments that were more effective at the start of the experiment. The cumulative difference in mite drop between the oxalic acid and oxalic acid plus adjuvant treatment with the solvent control are likely due to increased mite drop compared to the control up to day 9, as the samples from days 14-16 and 21-23 indicated decreased levels of mite drop. This is most likely due to the fact that oxalic acid is not effective at controlling mites under brood cappings (38,39). It is also possible that oxalic acid extended-release formulations using glycerin and Swedish sponges do not have longevity to control *Varroa* over 14 days. While studies have been performed on similar oxalic acid extended-release treatment for *Varroa* control, no currently published studies have addressed the persistence of oxalic acid in the treatments beyond taking measurements of mite levels in colonies.

The treatment time of 23 and 22 days for year 1 and year 2, respectively, were used for logistical reasons, but typical extended-release products for *Varroa* control are labeled for 42 to 56 days. Residue testing on treatment strips would be useful in determining if treatments are effective for a full 42- to 56-day treatment period. Comparisons of oxalic acid spreading throughout the colony at different time points (83) would also be useful in determining the rate at which the treatment can take effect. Further testing is required to determine the optimal oxalic acid concentration, the optimal adjuvant concentration, and whether glycerin alone or a glycerin-water mixture should be used. While treatments were all performed in 3 geographically distinct apiaries, efficacy for many *Varroa* control pesticides can vary with climate (84–88). Further testing of this formulation should be performed in other climatic regions.

Adjuvants are underutilized for pesticide formulations to control honey bee pests. While adjuvants are valued globally at 3.8 billion USD in 2023 (89), we could find no peer-reviewed journal articles that sought to improve control of pesticides used within honey bee colonies making use of adjuvants. Five adjuvant products, including two polyethylene/polypropylene glycol ethoxylates (Ecostep AE-13® and BC-12®), two fatty acid ethoxylates (Ecostep CE-13® and SE-11®), and one organo-silicone (Silwet L-7500 Copolymer), were investigated in this study, but there are many other types of adjuvants that may improve efficacy of *Varroa* control applications, including other non-ionic surfactants, crop oils, seed oils, hydrocolloid polymers, or combinations (90). Formulation of *Varroa* control products could take advantage of the spectrum of adjuvants to enhance new active ingredients for *Varroa* control (48,91).

All five adjuvants used in this study claim to act as a wetting agent by reducing surface tension of the pesticide mixture. These adjuvants are designed to be used with a water solvent but are soluble in glycerin. Other homemade oxalic acid extended-release formulations have used a mixture of glycerin and water (88,92), which may change the activity of an adjuvant. The adjuvant in the formulation may improve the efficacy of the miticide through increased spreading on the honey bees or across the *Varroa* themselves, thereby increasing the exposure of *Varroa* to the miticide. The mechanisms of increased spreading of the pesticide throughout the colony and across bee cuticle is critical for the exposure of *Varroa* to the miticide, as most *Varroa* on adult bees are located under the honey bee abdominal sternites (93). Adjuvants may act through mechanisms that improve spiracular or cuticular penetration of pesticide in *Varroa*, as demonstrated in other arthropod pests (57,94–97). Adjuvants may also have intrinsic toxicity to *Varroa*, as demonstrated in spider mites (97–100), aphids (100–103), thrips (104), cockroaches (105), and mosquitoes (51,106). Other adjuvants not used in this study have also demonstrated toxicity to honey bees (53,107–112) and other bee species (113,114), so it is important to perform preliminary testing prior to including adjuvants in candidate miticide formulations. The mechanism of action for adjuvant toxicity to arthropods is not well understood, but it is thought that adjuvants act similarly to insecticidal soaps (53,99,115,116), which can disrupt the arthropod cuticle, break down cell membranes, and reduce water surface tension to cause spiracular drowning (106,117).

## Conclusion

Field trials with oxalic acid and adjuvant in glycerin caused a significant decrease in *Varroa* in year 1 and a significant reduction in *Varroa*, relative to the solvent control, in year 2. On the other hand, field trials with oxalic acid alone did not reduce *Varroa* loads in colonies relative to the solvent control. While, oxalic acid extended-release formulations in glycerin have been found to be effective in some studies (16,92,118,119), other studies found them to be ineffective (88). The field trials reported in this study show that oxalic alone did not provide effective *Varroa* control. Additionally, mite drop data indicate increased speed for the miticidal effect when an adjuvant is included with oxalic acid. Adjuvants with low-risk to honey bees can improve miticidal active ingredients to reduce levels of *Varroa* in colonies. Oxalic acid extended-release products utilizing adjuvants should be further developed and registered for use through the US EPA or appropriate regulatory agency prior to use by beekeepers. There is a need for improved strategies for managing *Varroa* among beekeepers. Improving formulation chemistry is a critical tool to improve efficacy of chemical controls that can be incorporated within an integrated pest management program (20,120).

## Acknowledgements

We thank Brooke Fries and Emily Greenland for their contribution to methods development. We thank Lauren Tarver for her contributions to acute toxicity testing, cage manufacturing, setup of laboratory assays, collection of sticky boards, and sticky board *Varroa* counts. We thank Makayla Phillips for her contributions to setup of laboratory assays, setup of field trials, collection of sticky boards, and counting *Varroa* in sticky boards and ethanol washes. We thank Frank Rinkevich for advice on cage test methodology. We thank Stepan for providing the Ecostep adjuvant samples and Momentive for providing the Silwet adjuvant sample.

## Supplemental Information

### S1 Text. Supplemental Tables and Figures

**S1 Table A.** 24-hour bee mortality and *Varroa* mite drop efficacy for cage trials, expressed as a percentage, with range of standard error listed in parenthesis. The P-value of each treatment compared to the glycerin control and active ingredient control were determined via a pairwise Wilcoxon Rank-Sum Test using Benjamini-Hochberg post-hoc correction. Statistically significant differences (P < 0.05) are indicated with an asterisk (*).

**S1 Table B.** Year 1 and 2 ethanol wash data for each treatment, where rate is defined as *Varroa* infestation rate per 100 bees. The value listed in parenthesis for the initial and final rate mean indicates standard deviation; the range listed in parenthesis for the mean difference indicates standard error. The column “n” indicates the number of colonies used for analysis for each treatment. P-value within treatments was determined using a one-sided t-test on the difference within treatments and P-value between treatments and each of the controls was determined with an ANOVA with Tukey’s Post-Hoc test of the change in mite levels, where an asterisk (*) indicates a statistically significant test (P < 0.05).

**S1 Figure A.** Cage design for laboratory cage trials, with approximately 300 bees added to each cage.

**S1 Figure B.** *Varroa* levels from pre- and post-treatment alcohol washes in the year 1 (A) and year 2 (B) field trials. Points indicate mites per 100 bees for each colony, and lines connect paired pre- and post-treatment data for each individual colony. Significant difference within treatments, indicated by an asterisk, was determined for the Oxalic Acid Plus Adjuvant Treatment in Year 1 (P = 0.0122, n=6, t= –3.189).

### S2 File. R Code for Mixed Model for Repeated Measures (MMRM) analysis of mite drop

**S3 Dataset. Adjuvant Toxicity Testing Dataset.** Adjuvant dose is listed as a percent composition by volume. Insecticide dose is listed as multiples of the application rate (X), where 1X Mustang Maxx is equivalent to 0.0186 percent composition by volume of formulation, or 0.00170 percent composition by weight of zeta-cypermethrin active ingredient.

**S4 Dataset. Cage Trial Dataset.**

**S5 Dataset. Year 1 Field Trial Dataset. S6 Dataset. Year 2 Field Trial Dataset.**

